# Rad5 and Telomere Replication

**DOI:** 10.1101/2025.01.16.633138

**Authors:** Stefano Mattarocci

**Affiliations:** Université Paris-Saclay, Université Paris-Cité, CEA, Inserm, Institut de biologie François Jacob, UMR Stabilité Génétique Cellules Souches et Radiations, Fontenay-aux-Roses, France

**Keywords:** DNA replication, DNA repair pathways, Rad5, telomere, cell fate, homologous recombination, ALT

## Abstract

The DNA Damage Tolerance pathway (DDT) is one of the major mechanisms for resolving replication fork blocks. A central factor for DDT is the fork-associated clamp, PCNA, which can undergo to mono- or polyubiquitination, leading to error-free or error-prone modes of repair, respectively. The Rad5 protein in the yeast *Saccharomyces cerevisiae* plays important roles in both pathways: promoting the error-free mode by PCNA polyubiquitination and interacting with specialized DNA polymerases involved in the error-prone pathway. Rad5 also associates with telomeres, the repetitive DNA regions present at the ends of chromosomes. Telomeric DNA, tightly bound by tandem proteins arrays, poses unique challenges to replication fork progression. Here, I review the current understanding of the link between Rad5 and telomeres and present evidence that Rad5 is associated with yeast telomeres, with notable enrichment during telomere replication. This data highlights a connection between telomeres and a key DDT factor in unperturbed wild-type cells, raising intriguing possibilities about the functional interplay between telomere replication and DNA damage tolerance mechanisms.

## Rad5: a central factor for the DDT pathway

The DNA replication process ensures that the genome is fully duplicated prior to cell division. However, replication can be challenged by a multitude of endogenous and exogenous stresses. Endogenous stresses arise from natural barriers within the DNA sequence, such as tightly packed chromatin, repetitive sequences, DNA secondary structures, or DNA damages caused by reactive oxygen species (Saxena & Zou, 2022). These obstacles can impede replication fork progression (Saxena & Zou, 2022). Similarly, exogenous factors, such as exposure to UV or chemical agents, induce DNA lesions that, if not repaired, can also challenge replication (Kciuk et al., 2020). Both endogenous and exogenous stresses can lead to stalled replication forks. Unresolved stalled forks can result in the collapse of the replication machinery, resulting in DNA breakage and compromising overall genome stability. To counteract these threats, cells have evolved dedicated pathways, including the DNA damage tolerance pathway (DDT), also referred as post-replication repair pathway. The DDT pathway plays a role in bypassing DNA lesions that would otherwise block replicative fork progression, thereby preventing fork collapse and maintaining genome stability (Ulrich, 2011; Branzei & Szakal, 2016; Arbel et al., 2020, 2021). The key player of the DDT pathway is PCNA, a sliding homotrimeric clamp for DNA polymerases and an essential component of the replication machinery (Moldovan et al., 2007; Arbel et al., 2020, 2021; Bellí et al., 2022).

Stalled replicative forks frequently result in polymerase-helicase uncoupling, and thereby excessive production of single-stranded DNA (ssDNA), whose coverage by RPA can lead to an activation of the S-phase checkpoint (Liao et al., 2018; Técher & Pasero, 2021). In *Saccharomyces cerevisiae*, the RPA-ssDNA filament interacts with and engages Rad18, an E3 ubiquitin ligase, which in turn recruits Rad6, an E2 ubiquitin conjugase. The Rad18-Rad6 complex monoubiquitinates PCNA at lysine 164 (Liefshitz et al., 1998; Hoege et al., 2002). This step is crucial to allow the recruitment of translesion-synthesis (TLS) DNA polymerases (in budding yeast: Rev1, Pol ζ (Rev3-Rev7-Pol31-Pol32), Pol η (Rad30)). These error-prone polymerases are less hindered by DNA lesions, therefore allowing replication through the lesions (error-prone bypass pathway) (Prakash et al., 2005). On the other hand, PCNA_K164_ Ub (but not the unmodified PCNA) can be polyubiquitinated at lysine 63 by the E2 ubiquitin conjugase Ubc13-Mms2 and the E3 ubiquitin ligase Rad5 (Ulrich, 2001; Hoege et al., 2002; Torres-Ramos et al., 2002; Xu et al., 2015). This modification activates the error-free DDT pathway, which relies on a transient template-switching (TS) mechanism where the stalled DNA strand utilizes the newly synthesized sister chromatid strand as a template to bypass the lesion (Minca & Kowalski, 2010; Giannattasio et al., 2014).

Rad5 belongs to the SWI/SNF2 family of ATPases and contains seven conserved helicase-like motifs (Johnson et al., 1992, 1994; Ball et al., 2014). Rad5 contains also a RING finger domain, characteristic of the ubiquitin ligase enzymes, which is inserted between the helicase motifs III and IV (Lorick et al., 1999; Unk et al., 2010). Additionally, the N-terminal region of Rad5 harbors a HIRAN domain, proposed to function as DNA-binding domain (Iyer et al., 2006; Fan et al., 2018).

Rad5 interacts with Rad18 and PCNA, and both interactions are important for its localization at sites of DNA damage (Ulrich & Jentsch, 2000; Moldovan et al., 2007; Carlile et al., 2009). The greater UV-sensitivity of cells lacking Rad5 compared to cells lacking Mms2 or Ubc13 suggests that Rad5 could play additional roles beyond the activation of the TS pathway (Gangavarapu et al., 2006). Firstly, Rad5 has been shown to interact with Rev1, one of the TLS polymerases (Pagès et al., 2008; Kuang et al., 2013; X. Xu et al., 2016). Genetic evidence highlights the existence of two independent pathways acting when the TS pathway is suppressed. In these pathways, the TLS polymerases are recruited, either via the mono-ubiquitinated PCNA at lysine 164 or by Rad5, which in this case acts as a scaffold protein (Jiang et al., 2023). Therefore, Rad5 could play a role in the choice between error-prone and error-free DDT pathways (Pagès et al., 2008; X. Xu et al., 2016; Ortiz-Bazán et al., 2014; Choi et al., 2015; Gallo et al., 2019). Secondly, *in vitro* study indicates that Rad5 can bind various DNA structures *in vitro* (Blastyák et al., 2007) and possesses DNA helicase activity specific for replication fork reversal (Shin et al., 2018). Both Rad5 and its human orthologue, *HLTF*, can regress replication fork to form “chicken foot” structures, which are thought to be important for the stabilization and the rescue of stalled replication forks (Shin et al., 2018; Bai et al., 2020). Fork reversal can rescue DNA replication after a block by promoting the DDT pathway or by limiting the uncoupling of leading and lagging DNA strand synthesis, thereby preventing the accumulation of single-stranded DNA (Neelsen & Lopes, 2015; Berti & Vindigni, 2016). Overall, these data suggests that Rad5 contributes to DNA lesion bypass through multiple pathways: strand invasion-dependent template switching (TS), the promotion of TLS polymerase activity and replication fork reversal.

### Telomeres: a challenging region for DNA replication

Specific genomic regions can challenge the DNA replication machinery. Among these, telomeres - the nucleoprotein structures capping the ends of chromosomes - pose problems for the replication process even in absence of exogenous replication stress (Mason-Osann et al., 2019; Cicconi & Chang, 2020; Stroik & Hendrickson, 2020; Bonnell et al., 2021). Telomeres are composed of G-rich repetitive sequences tightly bound in tandem by specialized proteins that protect the terminal portions of chromosomes (De Lange, 2018). These G-rich sequences could form secondary structures, such as G-quadruplex (Traczyk et al., 2021; Xu & Komiyama, 2023), which are known to impede replication fork progression (Joo et al., 2024). Stabilizing these structures with chemical compounds reduce cell viability, possibly due to telomere replication failure (Vertecchi et al., 2022). Moreover, telomeres are transcribed into non-coding RNA known as TERRA, which could form R-loops, *i.e*. DNA-RNA hybrids that could impede fork progression (Rivosecchi et al., 2024; In et al., 2025).

Another important source of replication obstacles at telomeres is represented by the proteins that directly bind to the telomeric repeats: Rap1 in budding yeast, Taz1 in fission yeast and TRF1-TRF2 in mammals (König et al., 1996; Cooper et al., 1997; Kupiec, 2014; De Lange, 2018). Despite structural differences, Rap1, Taz1 and TRF1/2 utilize conserved Myb-like motifs to bind telomeric DNA (König et al., 1996; Cooper et al., 1997; Matot et al., 2012; Chen, 2019). In *S. cerevisiae*, Rap1’s tight binding to telomeric repeats (Williams et al., 2010; Analikwu et al., 2023) represents a significant block to replication fork progression (Douglas & Diffley, 2021). Rap1 also impede other DNA transactions, including transcription (Wu et al., 2018; Challal et al., 2018) and condensin loop extrusion (Analikwu et al., 2023). While tight DNA-protein binding is essential for efficient telomere end protection (Mattarocci et al., 2024), it may come at a cost, creating challenges for specific DNA transactions, including DNA replication.

Several evidence point out that telomere regions could be “hot-spots” for the accumulation of replication intermediates, which in turn recruit various DNA repair factors, including checkpoint proteins, helicases and the Rad51 recombinase (Brenner & Nandakumar, 2022; Maestroni et al., 2017; Higa et al., 2017). Several proteins help the replication machinery in passing through telomeres (e.g.: the Pif1 and Rrm3 helicases) (Makovets et al., 2004; Geronimo & Zakian, 2016). Telomerase - the holoenzyme elongating short telomeres - can also be considered a repair factor, since it repairs broken telomeres resulting from replication defects by extending them (Matmati et al., 2020).

In the absence of telomerase, other compensatory pathways become essential for telomere stability. As cells divide, telomeres progressively shorten, eventually to a point where they become dysfunctional and prone to fusion or extensive resection (Teixeira, 2013; Pobiega et al., 2021). This uncapping/de-protection of telomeres triggers a persistent Mec1^ATR^-dependent checkpoint response, ultimately leading to cell death, an outcome termed “telomere crisis” (Lundblad & Szostak, 1989). Critically short telomeres whose replication has been defective could be viewed as dysfunctional and in need to be repaired. Consequently, the survival and proliferation of telomerase-negative cells depends on the activation of repair pathways and thus of the recruitment of repair factors at chromosome ends. Rad5 has been proposed to be involved in this response.

### Does Rad5 play a role in telomere replication?

The link between Rad5 and telomere biology was discovered serendipitously. In budding yeast, it has been proposed that artificially shortening a single telomere accelerates the telomere crisis caused by the loss of telomerase (Abdallah et al., 2009). Notably, the W303 genetic background used in that initial study, carries a mutant allele of the *RAD5* gene, *rad5-535*. This allele harbors a point mutation in one of the seven consensus motifs of the Rad5 helicase domain (Fan et al., 1996). Cells with the *rad5-535* mutation exhibit defects in the error-prone TLS pathway, resulting in heightened sensitivity to DNA-damaging agents such as MMS and increased genomic instability (for more details, see *“Choose your yeast strain carefully: the RAD5 gene matters”* (Elserafy & El-Khamisy, 2018). Therefore, the reduced cell viability observed upon the shortening of a single telomere in this genetic background could have been partially attributed to the defective *rad5-535* allele, suggesting a potential role of Rad5 at telomeres. Subsequent work confirmed this hypothesis: Rad5 loss exacerbates cell death caused by an artificially short telomere in telomerase-negative cells (Fallet et al., 2014). Interestingly, another DDT pathway factor, Mms2, was also found to be involved in the maintenance of short telomeres in the absence of telomerase (Fallet et al., 2014). The discovery that Rad5 and the DDT pathway have a role at telomeres is intriguing, as it directly links telomere replication to factors that assist DNA replication under stress. Supporting this idea, Rad5 binds telomeres in telomerase-positive cells (Fallet et al., 2014). Despite these initial hints, direct evidence of Rad5’s implication in telomere replication remained unclear. Thus, I investigated whether Rad5 could be enriched at telomeres during their replication.

In budding yeast, there is a temporal program for DNA replication, where specific origins fire early in S phase (early origins) and others later (late origins). Telomeres are among the last regions to replicate (McCarroll & Fangman, 1988; Raghuraman et al., 2001; Theulot et al., 2025). To investigate whether Rad5 associates with replicating telomeres, I used Chromatin Immunoprecipitation (ChIP) coupled with cell cycle synchronization during S phase (Bianchi & Shore, 2007; Mattarocci et al., 2014; Hafner et al., 2018). Telomerase-positive cells were arrested in G1 and subsequently released at a low temperature (18.5°C) to slow down the replication fork progression, thereby improving the temporal resolution between early and late origins.

To track replication fork progression, I monitored the recruitment of the single-stranded DNA binding protein RPA (Waga & Stillman, 1998; Mattarocci et al., 2014; Hafner et al., 2018). As shown in **Figure 1A & B** (top graphs) and **Suppl. Fig. 1**, and consistently with previous reports (Mattarocci et al., 2014; Hafner et al., 2018), RPA was recruited to the early origin *ARS607* approximately 45 minutes after release from the G1 block, producing a sharp ChIP peak that reflect the high firing efficiency of early origins (Theulot et al., 2025). The mid/late origin *ARS522* was replicated predominantly at 60 minutes, with a broader peak indicative of its lower firing probability. As expected (Bianchi & Shore, 2007; Mattarocci et al., 2014), replication fork passage at two native telomeres (TEL VI-R and XV-L) occurred later, peaking between 75 and 90 minutes after release. RPA enrichment at telomeres was also higher throughout the cell cycle compared to its enrichment at internal origins. This elevated signal may reflect either RPA constitutive presence at telomeres or a higher background signal inherent to telomere regions in ChIP experiments.

**Figure 1.**
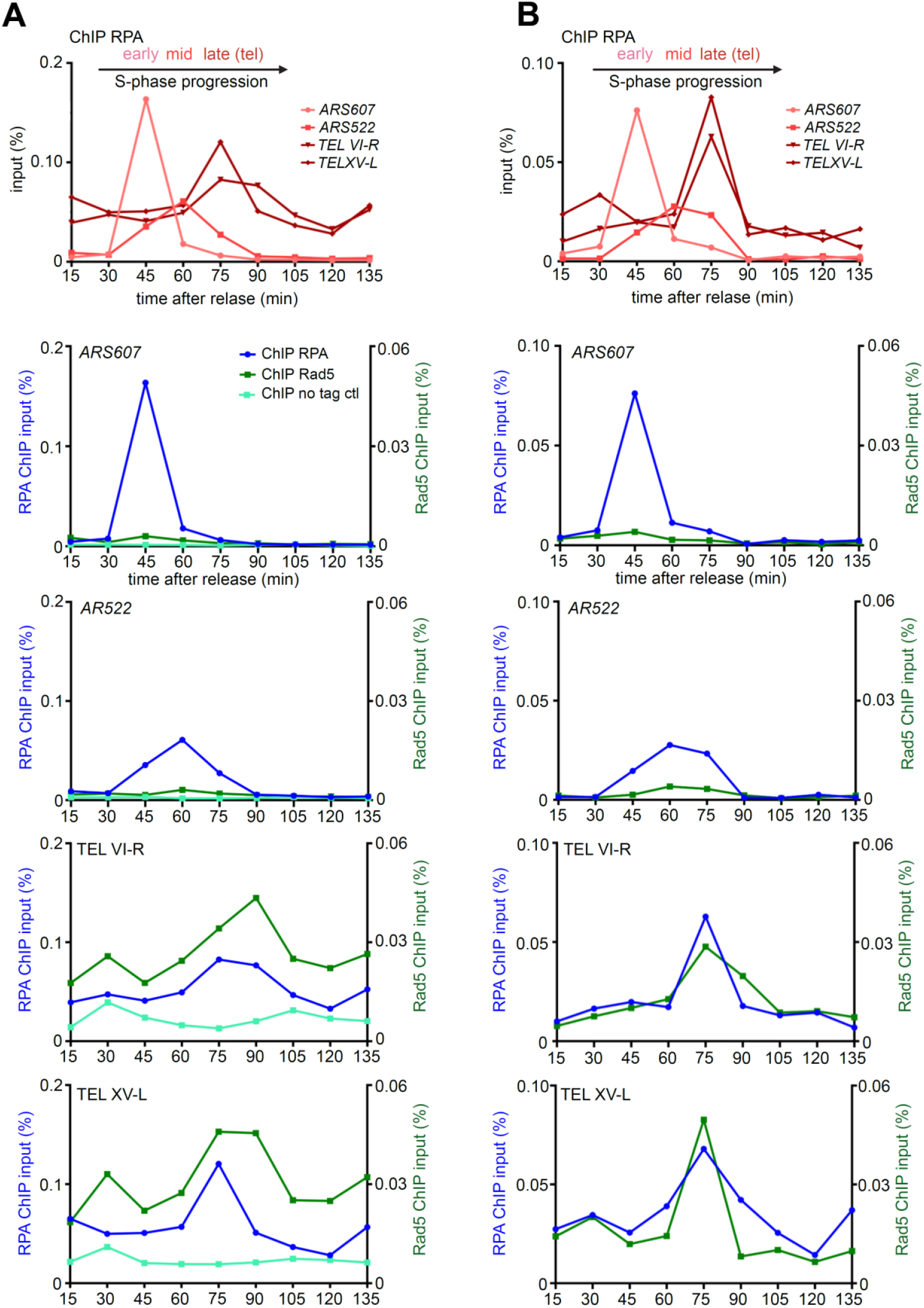
Rad5 is enriched at replicating telomeres. **A-B (upper graphs)**. Two biological replicates of RPA ChIP-qPCR experiments performed in Rad5-Myc tagged cells. Cells were blocked in G1 by the addition of α-factor and subsequently released at 18.5°C by washing away and degrading the α-factor by Pronase addition. Following release from the G1 block, samples were collected at 15-minute intervals for ChIP analysis (time point 0 min corresponds to Pronase addition). RPA enrichment at specific genomic loci was assessed: *ARS607* (early replicative origin, in light red), *ARS522* (mid/late replicative origin, in red), and subtelomeric regions TEL VI-R and TEL XV-L (in dark red, qPCR edge approximately 100 bp from the telomeric TG_1-3_ repeat). Results are shown as a percentage of the input fraction. **A-B (bottom graphs)**. Half of the samples processed for RPA ChIP (upper panels) were used for ChIP of Rad5-Myc (two replicates shown). To facilitate the comparison, RPA enrichment curves (blue) from the upper panel are also displayed in the lower panels. In the left replicate (A), a no-tag control ChIP (light green curve) is included, using anti-Myc immunoprecipitation in untagged *RAD5* cells. The background signal detected at native telomeres in the untagged strain was relatively high throughout the cell cycle compared to non-telomeric loci but remained approximately 3-fold lower than the Rad5-Myc signal and did not increase at the time of telomere replication (75-90 min).

The same ChIP samples analyzed for RPA recruitment were then assessed for the recruitment of Rad5, tagged with Myc epitopes (**Figure 1A & B**, bottom graphs; **Suppl. Fig. 1**). As a control, an untagged strain was included (**Figure 1A**). At the two internal origins, *ARS607* and *AR522*, a slight Rad5 specific enrichment above the control at the time of their respective replication (45 minutes and 60-75 minutes respectively) was detected. This suggests that Rad5 may associate with the replication forks in a subset of cells. In contrast, at the two native telomeres (TEL VI-R and XV-L), Rad5 enrichment was consistently higher throughout the cell cycle compared to the internal regions. Interestingly, Rad5 enrichment further increased during telomere replication (75-90 minutes), coinciding with the peak of RPA recruitment (**Figure 1A & B; Suppl. Fig. 1**). Overall, these findings suggest that Rad5 is constitutively present at telomeres and becomes further enriched during telomere replication in wild-type cells.

### Putative functions of Rad5 at telomeres

Here is described a connection between the replication of telomeres, a genomic region presumed to be inherently difficult to replicate, and Rad5, a protein helping DNA replication under stress conditions. It suggests that Rad5 is present at telomeres, possibly assisting replication the fork progression. It would be interesting to understand how Rad5 is recruited at telomeres. Recruitment could occur through interaction by other factors, such as proteins involved in DDT or telomere proteins, or through interaction between Rad5 HIRAN domain and DNA. Genetic analysis will be helpful in dissecting this recruitment mechanism. In addition, the functions mediated by Rad5 at telomeres in telomerase-positive cells remains to be explored. In telomerase-negative cells, Rad5 postpones telomere crisis, suggesting that Rad5-dependent template switching or fork reversal helps cells to cope with critically short telomeres (Fallet et al., 2014; Aguilera et al., 2020). Similarly, in telomerase-positive cells, Rad5 and template switching or fork reversal could help to rescue abnormally short telomeres that can arise sporadically even in the presence of telomerase. In this scenario, Rad5 could be constitutively recruited to telomeres in anticipation of these rare events. Alternatively, DDT pathways may operate at all telomeres, not only at the shortest ones. For instance, template-switch or fork reversal could help the recovery of replicative forks that pause at Rap1-bound telomere repeats. Further studies addressing the multiple functions of Rad5 at telomeres could provide key insights into its role in DDT and telomere replication.

## Materials and methods

### Yeast strains and primers used in this study

In this study, I used a *RAD5* corrected wild-type W303 *Saccharomyces cerevisiae* strain (*MATa bar1Δ ura3-1 trp1-1 leu2-3 112 his3-11 can1-100 RAD5 ADE2*), which served as the no-tag control for ChIP-qPCR experiments. The *RAD5* gene was tagged with 13 Myc epitopes using the *HIS3-MX6* selective marker (Longtine et al., 1998). The primers used for the qPCR step of the ChIP assay were as followed: TEL XV-L 5’-ATCGTGGTTCGCTGTGGTAT-3’ and 5’-AACCCTGTCCAACCTGTCTCC-3’; TEL VI-R 5’-TCCGAACTCAGTTACTATTGATGGAA-3’ and 5’-CGTATGCTAAAGTATATATTACTTCACTCCATT-3’; *ARS607* 5’-TCTGAACTGCAAATTTTTGTCATA-3’ and 5’-AGCCTTGTGCAGAAAGCATATGT-3’; *ARS522* 5’-CGTTCGAAAACCGGATATGT-3’ and 5’-CCCGATGACTACGAGGCTAT-3’; *OGG1* 5’-CAATGGTGTAGGCCCCAAAG-3’ and 5’-ACGATGCCATCCATGTGAAGT-3’.

### Cell synchronization

Cells were diluted to exponential phase in 540 ml of YPD medium and grown for 180 minutes at 30°C. To arrest cells in the G1 phase, α-factor (final concentration: 10^−8^ M) was added, and the cultures were incubated for an additional 100 minutes at 30°C. To release cells into S phase, they were washed three times with water and treated with 56 mg of nuclease-free Pronase (Millipore 537088R). The cultures were then incubated in 540 ml of YPD at either 18.5°C (**Figure 1**) or 19°C (**Suppl. Figure S1**). Cell-cycle synchrony and progression were monitored by microscopy.

### Chromatin Immunoprecipitation

ChIP was performed as previously described (Mattarocci et al., 2014; Hafner et al., 2018). For each ChIP culture, 40 ml of cells (10^7^ cells/ml) were fixed with 1% of formaldehyde for 5 minutes while rotating. The reaction was stopped by adding 2 ml of 2.5 M glycine. Fixed cells were then washed once with pre-chilled HBS buffer (50 mM HEPES pH 7.8, 150 mM NaCl) and frozen in 600 μl of ChIP lysis buffer with protease inhibitors (50mM HEPES pH 7.8, 150 mM NaCl, 1 mM EDTA, 1% NP40, 1% sodium deoxycholate). Cells were lysed using a Precellys homogenizer with zirconium/silica beads (BioSpec Products 11079105z). After bead removal, the lysate were pelleted, resuspended in 300 μl of ChIP lysis buffer, and sonicated using a Diagenod Bioruptor (high setting) for three cycles of 30 s ON + 30 s OFF in continuously cooled (4°C) water bath. The sonicated samples were clarified at maximum speed for 30 minutes at 4°C. 10 μl of the 300 μl total volume were put reserved as an input control (see below). Immunoprecipitation was performed by incubating the remaining lysate for 60 minutes at 4°C with agitation using either anti-Myc (Sigma-Aldrich 05-724) or anti-RPA (polyclonal Agrisera AS07214) antibodies. This was followed by a 120-minute incubation with magnetic beads crosslinked to sheep anti-mouse or anti-rabbit antibodies at 4°C with rotation. For the washing steps, the beads were collected using a magnet and washed as follows: first in 1 mL of ChIP lysis buffer, then transferred to a new tube and sequentially in the following buffers: (1) 50 mM HEPES pH 7.8, 150 mM NaCl, 1 mM EDTA, 0.03 % SDS; (2) 50 mM HEPES pH 7.8, 1 M NaCl, 1 mM EDTA; (3) 20 mM Tris pH 7.5, 250 mM LiCl, 1 mM EDTA, 0.5 % NP40, 0.5 % Sodium deoxycholate; (4) 20 mM Tris pH 7.5, 0.1 mM EDTA. Samples were eluted in 155 μl of elution buffer (buffer 4 + 1 % freshly added SDS) and reverse cross-linked in 140 μl of elution buffer at 65°C for 16 hours. The samples were then treated with proteinase K in TE buffer for 60 minutes at 37°C, followed by DNA purification using the DNA clean up kit (Thermo Scientific K0831). qPCR was performed using PowerSYBR green PCR master mix (Applied Biosystems 4367659) on an Applied Biosystems machine. The program used for amplification was: 95°C for 10 minutes, followed by 44 cycles of 95°C for 15 seconds and 60°C for 1 minute. The percentage relative to the input control (IN) was calculated using the following formula: % IP/IN = (E)^Ct(input)-Ct(IP)^, where E is in the PCR efficiency of the primers. The Ct_(input)_ values were corrected for dilution using the formula Ct’ = Ct – log_E_(30).

## Acknowledgements

I wish to thank Stephane Marcand for fruitful comments and his support; Florian Roisné-Hamelin, Karine Dubrana and Laurent Maloisel for helpful discussions and suggestions on the manuscript. During the preparation of this work, I used ChatGPT in order to enhance the fluency of the writing. After using this tool, I reviewed and edited the content as needed and take full responsibility for the content of the publication.

## Figure Legends

**Supplementary Figure 1.**
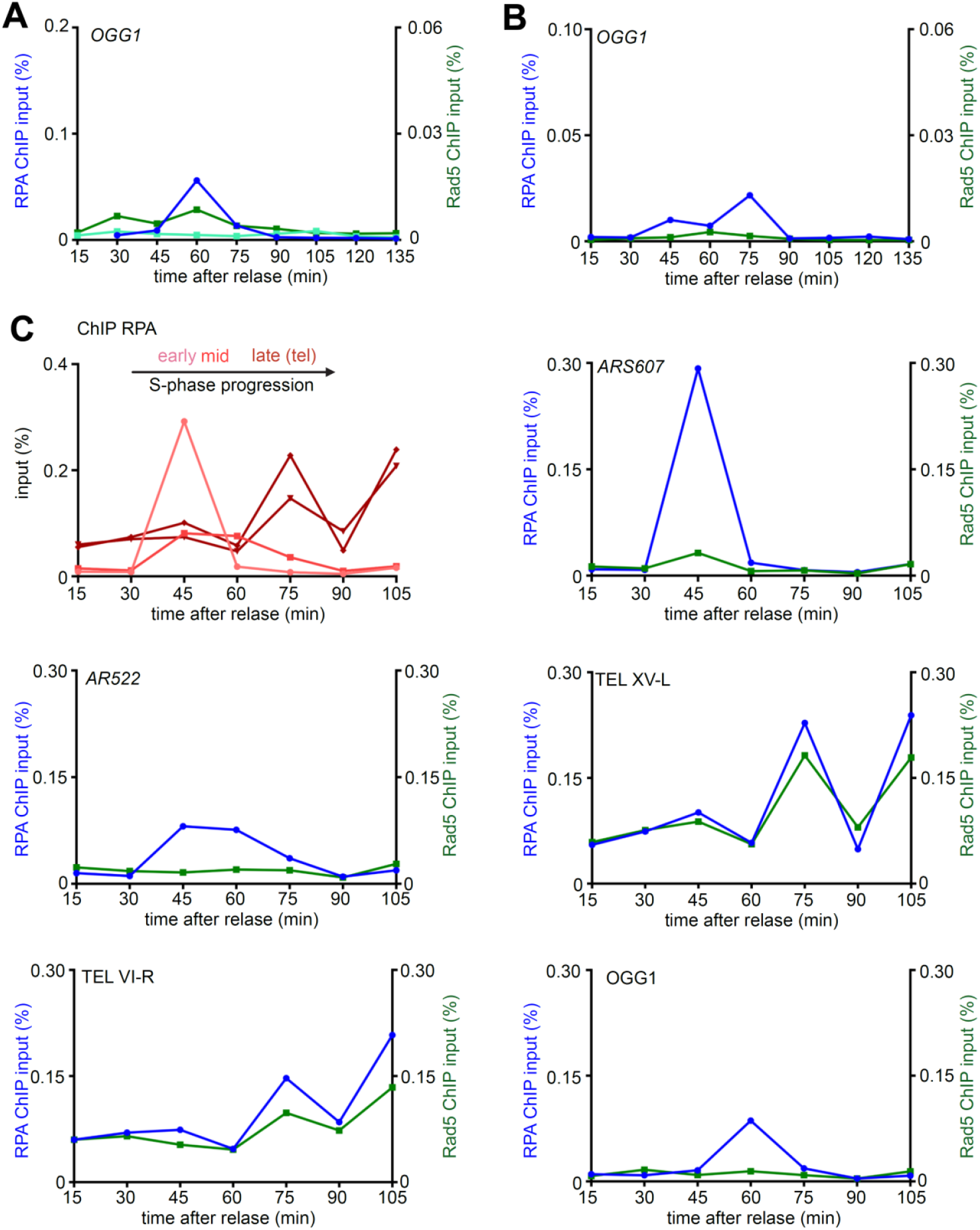
Rad5 is enriched at replicating telomeres. **(A-B)**. The ChIP samples used in **Figure 1** were analyzed to assess RPA and Rad5-Myc enrichment at the *OGG1* locus, an additional control for the Rad5-Myc ChIP. The replicative origin closest to the *OGG1* locus is the middle/late firing origin *ARS1307.5*. In the RPA ChIP, replicative fork passage is evident at 60 and 75 minutes. (**C)**. Triplicate experiment corresponding to **Figure 1**. In this replicate, cells were released into S phase at 19°C following Pronase addition.

